# Deep learning based root-soil segmentation from X-ray tomography images

**DOI:** 10.1101/071662

**Authors:** Clément Douarre, Richard Schielein, Carole Frindel, Stefan Gerth, David Rousseau

## Abstract

One of the most challenging computer vision problem in plant sciences is the segmentation of root and soil from X-ray tomography. So far, this has been addressed from classical image analysis methods. In this paper, we address this root/soil segmentation problem from X-ray tomography using a new deep learning classification technique. The robustness of this technique, tested for the first time on this plant science problem, is established with root/soil presenting a very low contrast in X-ray tomography. We also demonstrate the possibility to segment efficiently root from soil while learning on purely synthetic soil and root.

---

Machine learning is commonly used to designate the ensemble of techniques by which some information task (classification, quantification, prediction, identification) can be achieved automatically from a training data basis. So far, a vast majority of the work in plant phenotyping by imaging populations of plants has been developed in a classical image analysis method where the choice of features was made by humans. Some attempts at applying machine learning in plant sciences have recently appeared (see Ma et al. (2014) for a review). In these cases, however, the combination of the features was realized by the machine but the initial selection of the features was still mainly relying on human expertize. This expertize approach is possible when the phenotyping trait expected to be discriminant, for instance to identify clusters in the population of plants, is known before-hand. A computer assisted observation of fine traits overpassing human capacity of inference is nonetheless nowadays accessible. This even corresponds to a current huge trend in machine learning via the re-new of neural networks used under the form of the so-called deep-learning scheme LeCun et al. (1998, 2015) where the features on which the further information task is realized are directly learned by the computer. First applied in computer vision, deep learning is progressively being investigated for life science applications including biomedical imaging Greenspan et al. (2016), genetics Zhou and Troyanskaya (2015) and also very recently (so recently that it is even unpublished at the moment of the submission of this article) in plant phenotyping Pound et al. (2016). A limitation to the application of this technique to the field of plant phenotyping is that this requires huge training data sets to avoid overfitting. Such training data sets have to be annotated and at the moment only very few of these are available among the image analysis community interested in plant sciences (except for the important Arabidopsis case Minervini et al. (2015)). In this communication, we demonstrate how to circumvent this limitation and illustrate this on the soil/root segmentation problem.

The soil/root segmentation problem is one of the most challenging computer vision problem of plant sciences. Monitoring roots in soil is very important to quantitatively assess the development of the root system and its interaction with soil. This has been demonstrated to be accessible with X-ray computed tomography for single root systems Pound et al. (2013) and multiple root systems Mairhofer et al. (2015). The spatial resolution in X-ray computed tomography is very good with submillimetric resolution for entire root systems. However, the contrast between soil and root is rather limited Metzner et al. (2015). This makes this segmentation challenging. Currently there is only one soil-root segmentation software freely available for users Mairhofer et al. (2012). The process in Mairhofer et al. (2012) is initialized at the soil-root frontier manually by the expert. A level set method based on the Shannon-Jensen divergence of the gray levels between two consecutive slices is then applied to the stack of images slice by slice from the soil-root frontier down to the bottom of the pot where the root system is placed. The method is thus based on a prior of the human expert that the roots should follow some continuous trajectory along the root systems.

We propose another approach to address the root-soil segmentation from X-ray tomography, based on machine learning trained on annotated data sets. The communication is organized as follows. Useful machine learning concepts are first shortly recalled for non experts and in order to position our algorithm among the wide range of machine learning methods. Then, we go through our image segmentation algorithm which is applied in two experiments: In the first experiment, the root-soil image is simulated and has very weak contrast compared to the one used in Mairhofer et al. (2012) but in conditions which are however shown to be realistic. This is useful to stress the need for the development of additional tools to address in an extended range of soil-root contrast the problem of soil-root segmentation from X-ray computed tomography. The second experiment explores, on better contrasted root/soil, the possibility to realize efficient root-soil segmentation in real images after having trained our algorithm on purely synthetic data. We discuss in both cases the impact of the parameters of the proposed algorithm in terms of performance.

## 1. Concepts

### 1.1 Machine learning

Machine learning (ML) is a sub-field of artificial intelligence where algorithms learn from data and predict on data Bishop (2006). The most common task in ML is classification which consists in teaching the machine how to sort variables with certain attributes in one class or another. We shall now explain ML and further concepts from an image classification standpoint. In the image processing field, variables are images, and attributes are features. Extracted after filtering, these features carry information on the images which can be high or low level, local or global…For example, a feature “edge” could be the number of pixels considered as “edge”, by an edge-searching method, or their density, or another measure of their structure.

A typical example of classification in our case would be feeding the algorithm with images of cats and dogs and have it label each image as “cat” or “dog”. Classification is a supervised learning technique, which means we give the algorithm a batch of already labeled (=classified) images called a training set. The algorithm starts off by a learning phase, which means it is going to consider the features and the labels of the training images and search for the best way to use the former to match the latter. The second phase is the testing one, where the algorithm considers a new unlabeled set. Using the relationship between features and labels learned from the training set, if the training set was of a size large enough for the algorithm to learn on many different cases and “close enough” to the testing set, then the algorithm is capable of predicting the labels of the new images from their features. A large variety of classification algorithms exists, such as decision trees, k-nearest neighbors (KNN), support vector machine (SVM) …Bishop (2006).

### 1.2 Deep learning

Deep learning (DL) is a branch of ML imagined in the early 1980s, but its emergence had to wait for the computational power of the 2000s LeCun et al. (1998, 2015). It is a ML process structured on a so-called convolutional neural network (CNN). CNN works as a classifier, taking an image as an input and giving a label as an output.

A CNN is composed of several stacked layers of “neurons”, small computing elements, taking an input, passing it through a function and yielding an output. Neurons are connected to other neurons in upper or lower layers with each neuron link having a certain weight, and information flows through these connections. The input on the first layer of neurons is the image to be processed. Output of the first layer neurons is simply the values of the features computed on the image. These values then go through the network, undergoing subsampling, non linear transformation and linear combination as they pass through the layers, to finally yield a single output: the label of the image.

The CNN workflow is the following: During the training phase, features are computed for one image on the first layer, then flowed through the layers. A label is predicted: some neurons will have probably weighted in for one class, and some for the other. The predicted label is compared to the actual label. If the prediction is correct, then all neuron pathways which lead to this prediction are enhanced, *i.e.* they get more weight in the decision process next time, and all wrong neurons pathways are reduced. As training progresses, the best neurons are “selected” for the decision process.

The most interesting aspect of CNN is this: Each neuron can be seen as a feature extractor, applying a filter to the image. The filters from layers other than the first one are the result of a combination of filters from the layer above (because input of a neuron is simply a linear combination of outputs of neurons above). This has two profound consequences: (a) There is a hierarchy in the CNN neurons, *i.e.* neurons of the first layers detect low-level features (blobs, stripes, …), and this information passed on is used in the deeper layers to search for more complex features (spirals, faces, …); (b) since weights between neurons are constantly changing as training goes on, filters change too, and so computed features also change.

This last point is the crucial point that made CNN a revolution in ML algorithms: It implies that *features to use are found automatically during training*. Until CNN, the ML dogma was “training set + features to extract trained classifier”. Now, only the training set is necessary: ML has gone from feature engineering to feature learning.

### 1.3 Transfer learning

One of the problems of DL with CNN is that the learning phase, where the network undergoes weight modification, can be very time-consuming and may need a very large set of images.

In this article we use a common trick called transfer learning Pan and Yang (2010) to circumvent this problem. The idea is to use an already trained CNN to classify images. This trained CNN has been trained for a classification application which is not the one we want to perform. To understand why this approach for the classification is however working, let us recall that a CNN has two functionalities: (a) it has modeled features over training, (b) it classifies images given as input using these features. Only the (a) functionality of the pre-trained CNN is used in the transfer learning approach. This training-modeling phase of the CNN is realized beforehand on an existing very large database of images, manually classified in different cases among a large array of possible classes (e.g. dog races, guitar types, professions, plants, …). By being built on this large database, the CNN is assumed to have selected a very good feature space because it is now capable of sorting very diverse images in very diverse categories. The assumption is somehow grounded by the existence of common features (blobs, tubes, edges, …) in images from natural scenes. The study of these common features in natural scenes is a well-established problematic in computer vision which has been investigated for instances in gray level images Ruderman (1997); Gousseau and Roueff (2007), in color images Chapeau-Blondeau et al. (2009); Chauveau et al. (2010) and even in 3D images Chéné et al. (2013). It is thus likely that, since our images to be classified share some common features with the images of the database used for the training of the CNN, the selected features will also operate efficiently on our images. Please note that we do not use the (b) part of the pre-trained CNN because this CNN was trained on classes which are probably not the ones we are interested in, but then we simply feed our computed features to a classic classifier (as described in section 1.1) such as SVM.

## 2. Implementation

In this section we describe how, from the concepts shortly recalled in the previous section, we designed an implementation capable of addressing the soil-root segmentation from X-ray tomography images.

### 2.1 Application to image segmentation

We are interested in classifying each pixel of the image to segment as root or soil. However, the ML techniques presented in the previous section do not consider single scalars but images as input to realize a classification. Therefore, the idea is to consider for the classification of a pixel a small window, also called patch, centered on this pixel. Each patch is therefore a small part of the image, and its class (“soil/root”) is the class of the central pixel around which it was generated. It is these patches which are given to the pre-trained CNN. Once the features are computed for each patch, these features are given to a classic classifier. There is of course finally a training phase where this classifier receives labeled patches (coming from a segmented image), and then a testing phase where the classifier predicts the patch class.

### 2.2 Algorithm

Our algorithm, based on transfer learning and described as Algorithm 1, goes through three basic steps, both for training and testing: creating the patches around the pixels, extracting the features from these patches with a pre-trained CNN, and feeding them to a SVM. The training image is labeled (each pixel is labeled “part of the object” or “part of the background”), and these labels are fed to the SVM along with the computed features to train it. The trained SVM is then capable of predicting pixel labels from the testing image’s features. As underlined in Algorithm 1, the parameters to be tuned or chosen by the user rely on the size of the patch and the size of the training data set.

### 2.3 Material

In this article, we used an existing pre-trained CNN developed by Chatfield et al. (2014) and trained on Imagenet (http://www.vlfeat.org/matconvnet/pretrained/). We used this CNN because it is one of the most general ones (some CNN were trained on more specific cases such as face recognition), and so it can be expected to be more efficient in transfer learning on our problem. This network is composed of 22 layers and yields a total of 1000 features. Convolutional filters applied to the input image in the first layer of the CNN can be seen in Fig. 1. These filters appear very similar to wavelets Flandrin (1998) oriented in all possible directions. This is likely to enhance blob-like or tube-like structures such as the tubular roots or grainy blobs of the soil found in our X-ray tomography.

**Figure 1:**
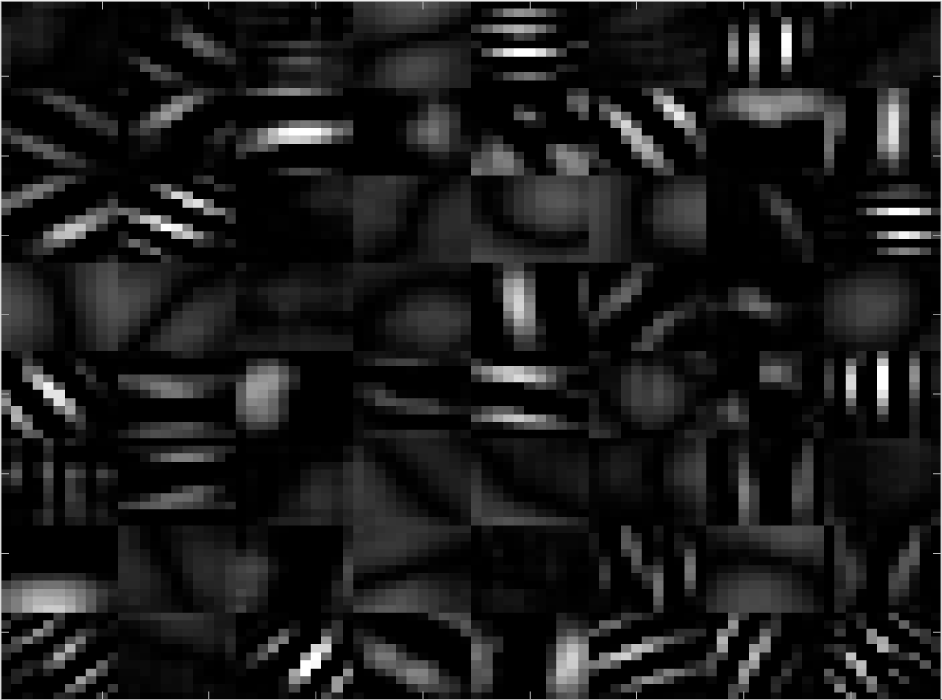
Convolutional filters selected by the first layer of the chosen CNN.

#### Algorithm 1 Proposed machine learning algorithm for image segmentation

~~~
1: CNN ← load(ImageNet.CNN);
2: training image ← root-soil-training-image.png; // image which segmentation we know
3: testing image ← root-soil-testing-image.png;
4: size images = size(training image);
5: nb of training pixels = **to be fixed by the user**; patch size = **to be fixed by the user**;
6:
7: // Training
8: training labels ← training image.labels;
9: training patches ← create patches(training image, nb of training pixels, patch size);
10: training features ← compute features(training patches, CNN);
11: trained SVM ← train SVM(training features, training labels);
12:
13: // Testing
14: testing patches ← create patches(testing image, size images, patch size);
15: testing features ← compute features(testing patches, CNN);
16: segmented image ← trained SVM.predict labels(testing features);
17:
18: **function** create patches(image, nb of pixels, patch size)
19:          **for** i=1:nb of pixels **do**
20:              *x ← random*; *y ← random*;
21:              patches(i) ← crop image(image, *x*, *y*, patch size); **return** patches;
22:
23: **function** compute features(patches,CNN)
24:    **for** i=1:length(patches) **do**
25:        features(i) ← CNN.compute features(patches(i)); **return** features;
~~~

The classifier used was a linear SVM. It was chosen after comparing performances on our data by cross validation with all other types of classifiers available in Matlab. Computing was run on Matlab R2016A, on a machine with an Intel (R) Xeon (R) 3.5 GHz processor, 32 GB RAM, and a FirePro W2100 AMD GPU.

All CT data are measured with an individually designed X-ray system at the Fraunhofer EZRT in Furth, Germany using a GE 225 MM2/ HP source Aerotech axis systems and the Meomed XEye 2020 Detector operating with a binned rectangular pixel size of 100*µm*. The source was operated at 175 kV acceleration voltage with a current of 4.7 mA. To further harden the spectra, a 1 mm thick copper pre-filtering was applied. The focus object distance was set to 725 mm and the focus detector distance to 827 mm resulting in a reconstructed voxel size of 88.9*µm*. To mimic the data quality typically occurring in high-throughput measurement modes, only 800 projections with 350 ms illumination time where recorded within the 360 degrees of rotation. This results in a measurement time of only 5 minutes for scanning the whole field of view of about 20 cm. The pot used for the measurement was a PVC tube with 9 cm diameter and only a small partial volume in the middle part of the whole reconstructed volume was used to reduce the segmentation time. The roots used as reference in experiment of section 3 and as experimental data for experiment of section 4 were maize plants of the type B73. During the growth period, the plants were stored in a Conviron A1000PG growth chamber. The temperature within the 12 hours of light is 21^°^ C and 18 ° C during the night. Two different soils were used for the two experiments. In experiment of section 3, the soils were the commercially available Vulkasoil 0/0,14 obtained from VulaTec in Germany. In experiment of section 4, the soils were the agricultural soil used in Metzner et al. (2015). Both soils were mainly mineral soils with a coarse particle size distribution. While the Vulkasoil inhibits in experiment of section 3 a very low contrast to the root system, the high amount of sand in the agricultural soil of section 4 sample increases the contrast visibly.

## 3. Segmentation of simulated roots

In this section, we designed a numerical experiment dedicated to the segmentation of simulated root systems after learning from other simulated root systems. Learning and testing images are both generated the same way:3D (244*244*26) soil images are real soil images coming from X-ray tomography. Root structure is generated from the L-system simulator of Leitner et al. (2010) under the form of Benoit et al. (2014a) presented in 3D in Benoit et al. (2014b). Simulated roots are added to the soil by replacing soil pixels by root pixels. The intensities of the roots are measured from a manual segmentation in real tomography images of maize in Vulkasoil. The estimated mean and standard deviation are given in Table 1. We then simulated with same mean and standard deviation for the gray levels the roots from a spatially independent and identically distributed Gaussian noise. As visible in Fig. 2, learning and testing images are not performed on the same realization of soil, nor the same root structure. This experiment is interesting because the use of simulated roots enable us to experiment various levels of contrast between soil and root. Also, since the L-system used is a stochastic process, we have access to an unlimited size of training or testing data set. It is therefore possible with this simulation approach to investigate the sensibility of the machine learning algorithm of the previous section to the choice of the parameters (size of the patch, size of the training data sets, …).

**Figure 2:**
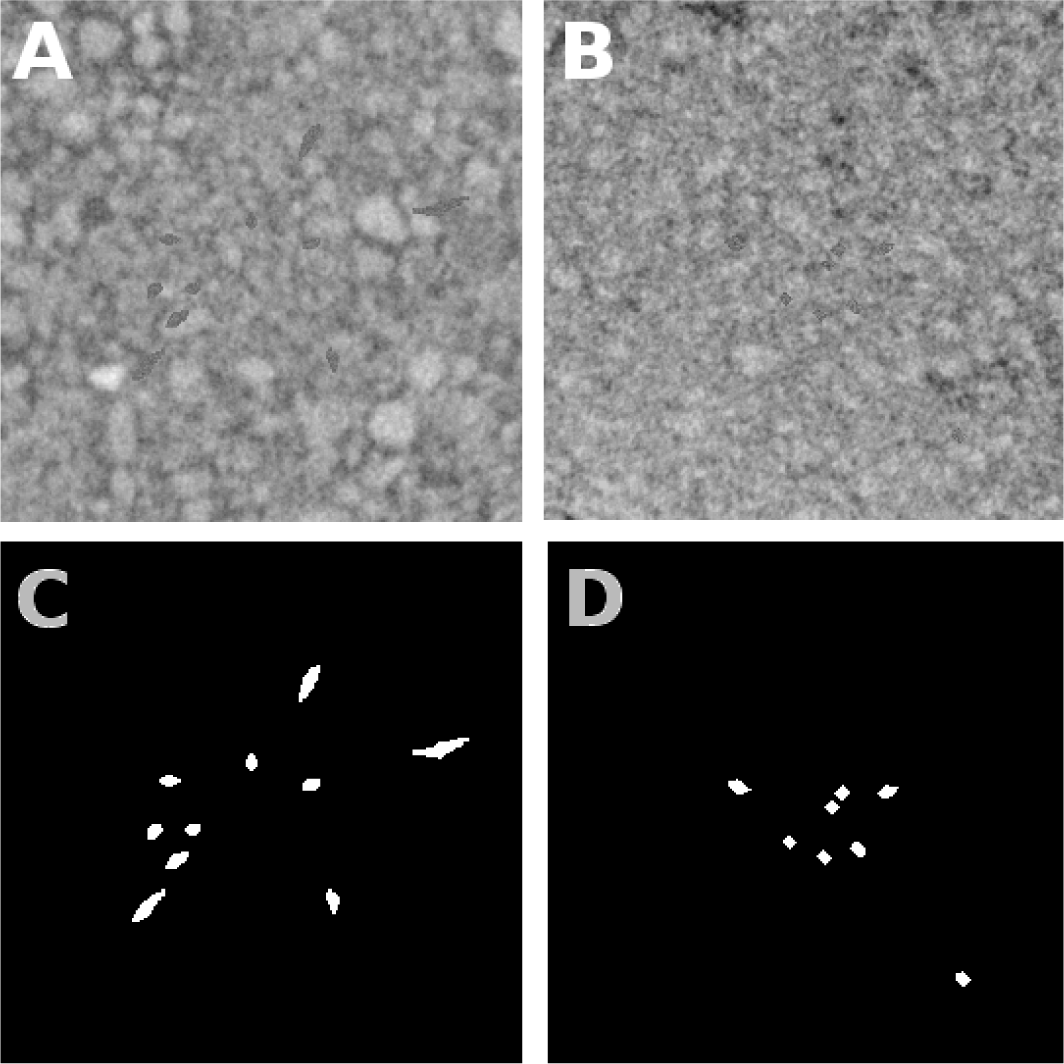
Presentation of simulated root systems. Panel A shows a slice of the training image (including soil and root). Roots were generated by simulating an L-system structure and replacing soil pixels by root pixels with gray intensity level drawn from a white Gaussian probability density function with fixed mean and standard deviation. Position of root pixels are shown in white in panel C which acts as a binary ground truth. Panel B gives a slice of the testing image where roots were generated the same way, and have the same mean and standard deviation than in panel A. Ground truth of Panel B is given in panel D.

**Table 1:** Mean (*µ*) and standard deviation (*ff*) values for the roots and soil in Fig. 2. Images are coded on 8 bits. Training and testing images have the same statistics.

| Statistics | Root $\mu$ | Root $\sigma$ | Soil $\mu$ | Soil $\sigma$ |
| --- | --- | --- | --- | --- |
| Values | 100 | 15 | 125 | 20 |

### 3.1 Nominal conditions

As nominal conditions for the root/soil contrast, we considered the first-order statistics (mean and standard deviation) of roots and soil given in Table 1 which corresponded to the contrast found in the real acquisition conditions of Maize in Vulkasoil (as described in section 2.3). As visible in Figs. 2 A and B, these are conditions which provide a very low contrast.

In the conditions of Table 1, with a combination of the information obtained with a patch of 2 pixels and 15 pixels (see section 3.2) and a training data set of 1000 patches, the segmentation results obtained are given in Fig. 3 with statistical performance given in the confusion matrix of Table 2. To summarize the performance of the segmentation with a simple scalar, we propose a quality measure *QM* obtained by multiplying sensitivity (which proportion of root pixels were detected as such) and specificity (which proportion of detected root pixels are truly root pixels)

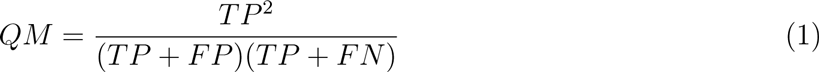

with *T P* being true positives, *F P* false positives and*F N* false negatives. The quality measure *QM* is maximized at 1 for perfect segmentation. For the segmentation of Fig. 3, *QM* = 0.23. As visible in Table 2 and in Fig. 3, the segmentation is not perfect especially since false positives outnumber true positives. However, the quality of the segmentation cannot be fully captured by sole pixel to pixel average metrics. The spatial positions of false positive pixels are also very important. And, as visible in Fig. 3, false positives (in yellow) are gathered just around the true positives and small false positives clusters are much smaller than the roots. This means that we get a good idea of where the roots actually stand and with very basic image processing techniques such as particle analysis and morphological erosion, one could easily yield a much better segmentation result. Also, when the segmentation is applied on the whole 3D stack of images, it appears in Fig. 3 panels E and F, that the overall structure of the root system is well captured by comparison with the 3D structure of the ground truth shown in Fig. 3 panels B and C. It is useful to recall, while inspecting Figs. 3 E and F that the process classification is realized in a pixel by pixel 2D process and it would also be possible to improve this result by considering 3D patches.

**Figure 3:**
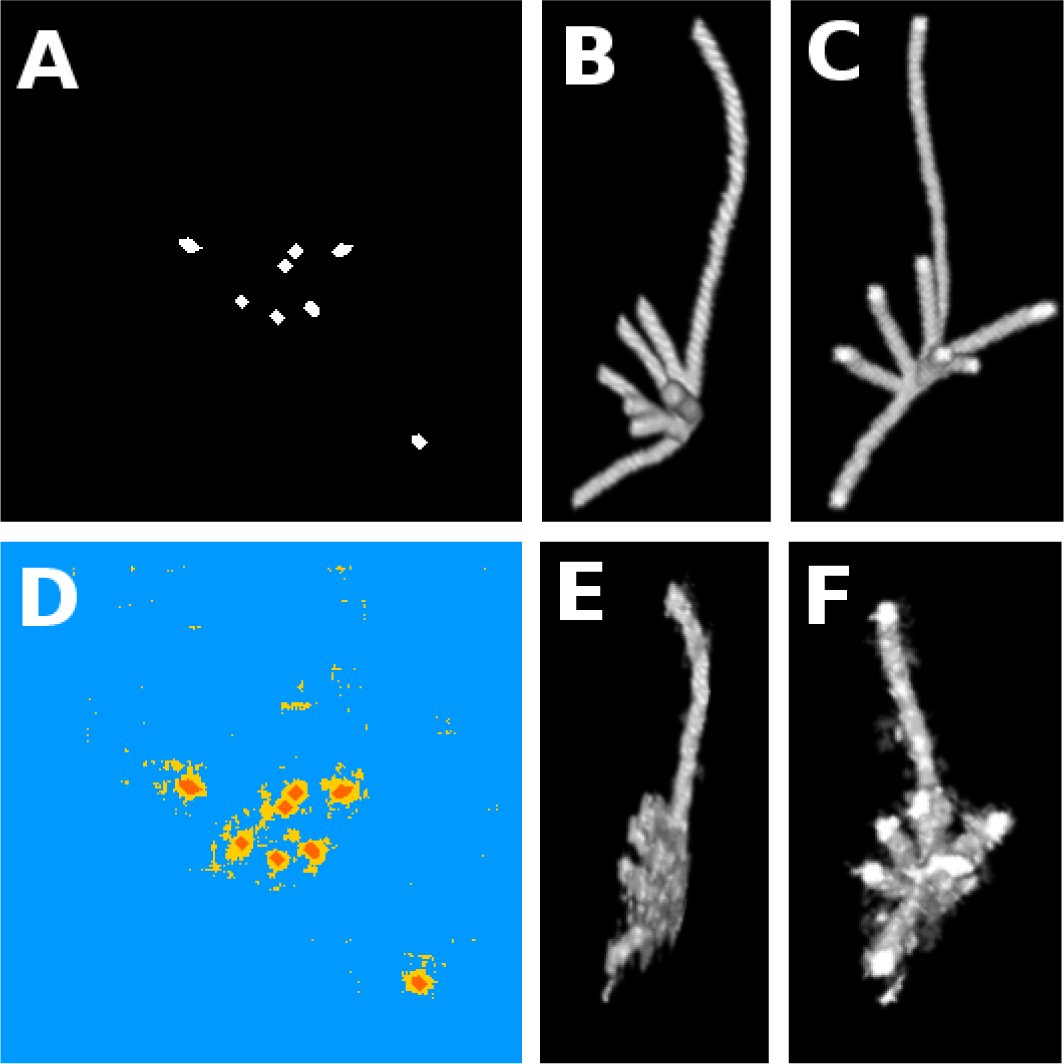
Experiments on simulated root. Panel A gives a slice of the binary ground truth (position of the roots in white). Panels B and C provide 3D view of ground truth from two different standpoints. Panel D gives the result of the segmentation. Blue pixels mean true negative (soil pixels predicted as such), yellow is false positive (soil pixels predicted as roots), orange true positive (root pixels predicted as such), and purple false negative (none in this result). Panels E and F are 3D views of the segmentation in the two viewing angles of panels B and C.

**Table 2:** Confusion matrix of results in nominal conditions as shown in Fig. 3. Total number of pixels was 1.784.744.

|  | Root pixel | Soil pixel |
| --- | --- | --- |
| Predicted root | 0.6% | 1.9% |
| Predicted soil | $3.10^{-3}\%$ | 97.5% |

To obtain the 3D (266*266*26 pixels) result of Fig. 3 E and F, about 2 hours were necessary. Computing time is mainly (95%) due to computing of the features on all the pixels of the testing image while the other steps had a negligible computation cost. Here we considered the full feature space (1000 features) from Chatfield et al. (2014). It would certainly be possible to investigate the possibility to reduce the dimension of this feature space while preserving the performance obtained in Fig. 3. Instead in this study, we investigate the robustness of our segmentation algorithm when the parameters or datasets depart from the nominal conditions exhibited in this section.

### 3.2 Robustness

A first important parameter for using our ML approach is the size of the learning data set. In usual studies based only on real data of finite size, the influence of the learning data set is difficult to study since increasing the learning data set necessitates to reduce the testing data set. With the data from the previous section, where roots are simulated, we do not have to cope with this limitation since we can generate an arbitrary large training data set. Figure 4 illustrates the quality of the segmentation obtained for different training data sets sizes on a single soil/root realization. A degradation of the results is visible only when the training size is decreased from 100 to 25 patches. Since the root simulator is a stochastic process, performance is also given in Fig. 5 as a function of the size of the training data set, in terms of box plot with average performance and standard deviation computed over 5 realizations for each size of training data set tested. As visible in Fig. 5, the average performance are found almost constant, and the increase of the training data set mainly benefits to the decrease of the dispersion of the results.

**Figure 4:**
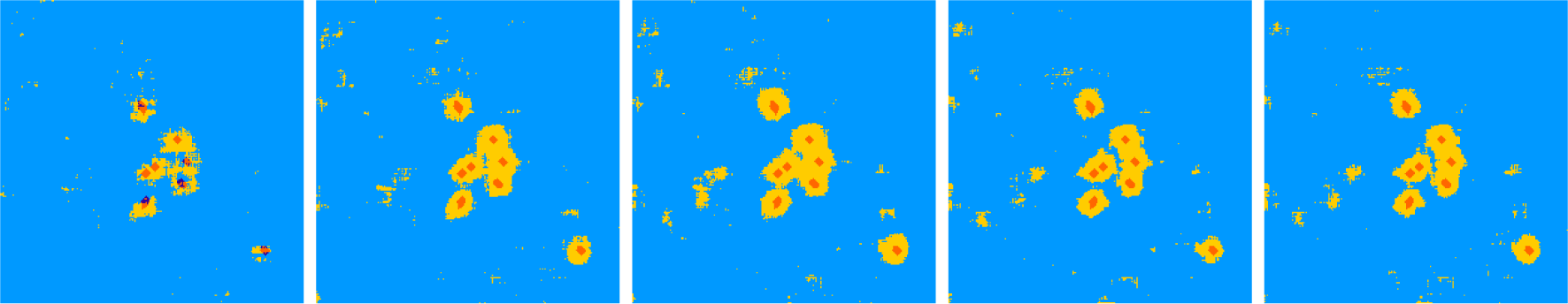
Segmentation results for a training size of 25, 100, 500, 1000 and 2000 patches, drawn from the 3D training image (see Fig. 2). The image to segment was of size 266*266 with 312 root pixels. Same color code as in Fig. 3.

**Figure 5:**
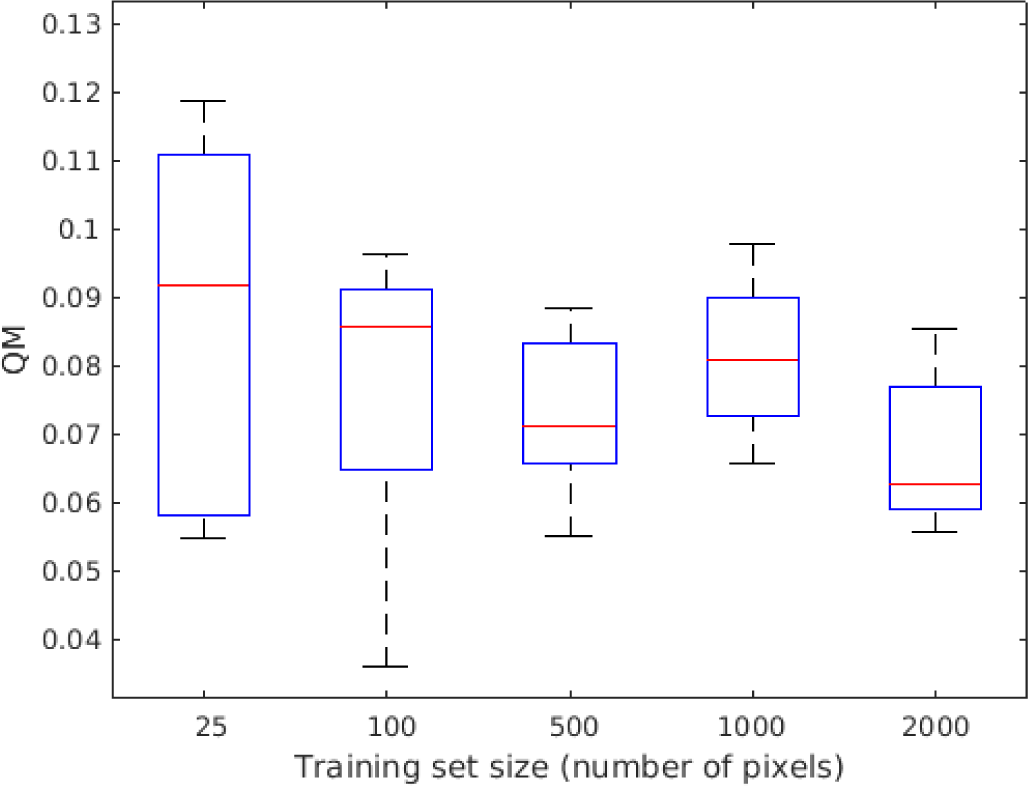
Quality of segmentation *QM* for a training size of 25, 100, 500, 1000 and 2000 patches. The root simulator being a stochastic process, the performance are given in terms of box plot with average performance (red line) standard deviation (solid line of the box) and max-min (the “mustach” of the box) computed over 5 realizations for each size of training data set tested.

A second parameter of importance to use our ML approach is the size of the patch. As visible in Fig. 6 left, decreasing the size of the patch produces finer segmentation of the roots and their surrounding tissue but also increases spurious false detection far from the roots. Increasing the patch size, see Fig. 6 right, produces a good segmentation of roots, with very few false detection far from the roots but with over segmentation on the tissue directly surrounding the root. An interesting approach can consist in combining the results produced from a small and a large patch by a simple logical AND operation which detects as root only the pixels detected as root for both sizes of patches. This was the strategy adopted in Fig. 3 which removes false detection far from the root while preserving a fine detection of the root with few false detection in the tissue surrounding the roots.

**Figure 6:**
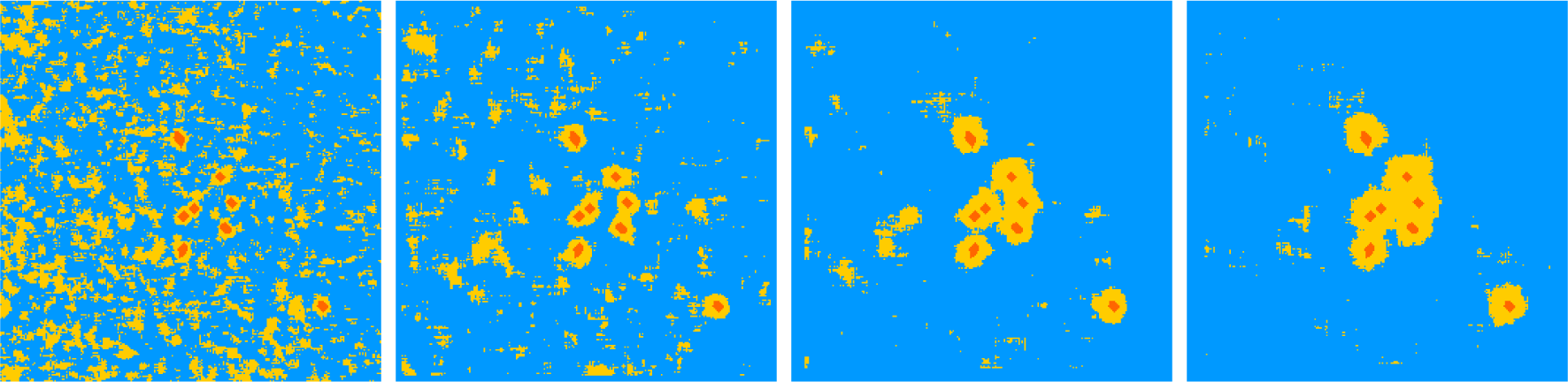
From left to right segmentation results for a patch size of 5, 15, 25 and 31 pixels wide. Roots in the image to segment could have a diameter of 10 to 15 pixels. Same color code as in Fig. 3.

## 4. Segmentation on real roots

In this section, we investigate the performance of our ML algorithm when applied to real roots. The contrast considered is higher than in the previous section (see detail in section 2) and corresponds to the one found in Mairhofer et al. (2012). We conducted the numerical experiment described in Fig. 7 designed for the segmentation of real root systems after learning from simulated root systems and simulated soil.

**Figure 7:**
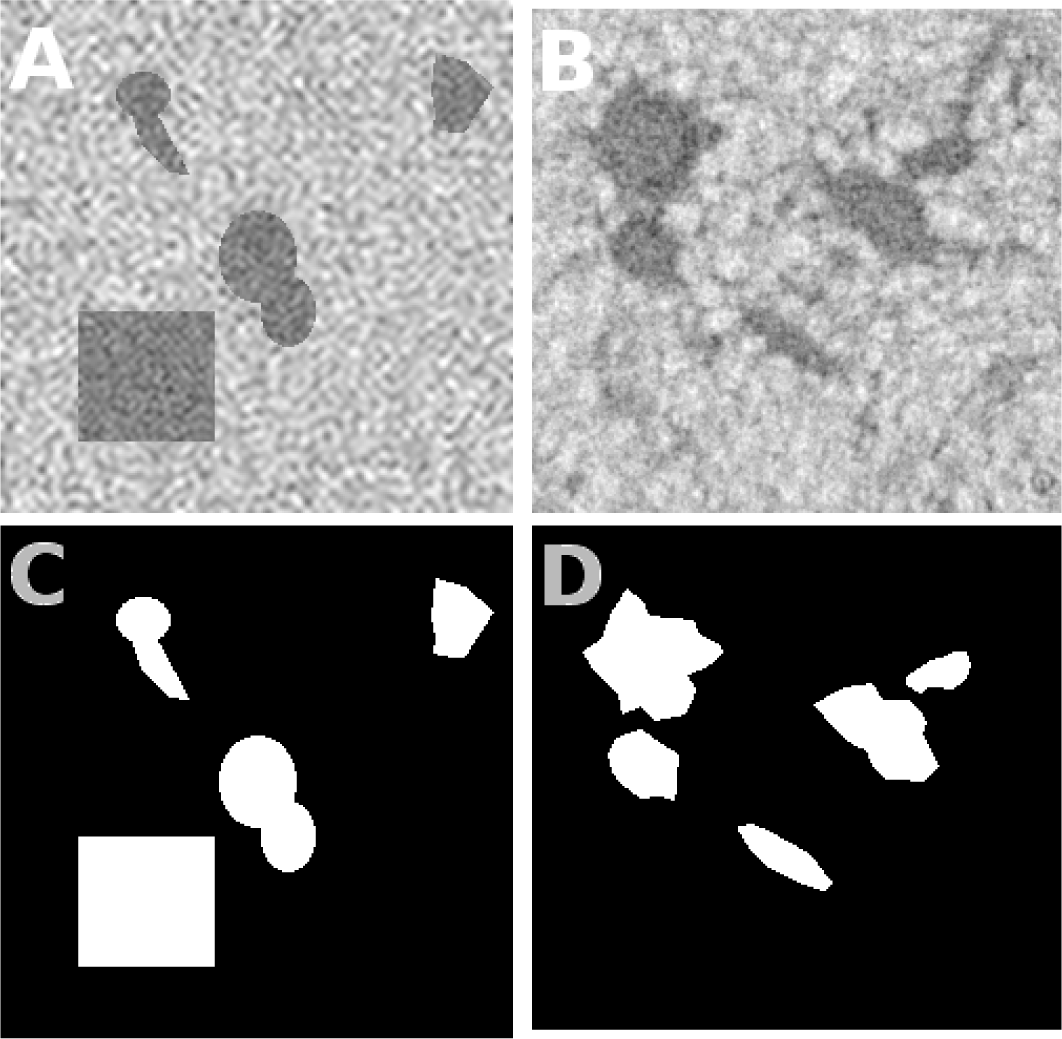
Experiment on real root: Panel A shows a slice of the training image (simulated). The corresponding binary ground truth is given in white in panel C. Root intensity values are generated from a white Gaussian probability density function with fixed mean and standard deviation, and are then low-pass filtered (see Figure 8) until the autocorrelation of the root (A) is similar to the one of real roots (B). Same goes for the soil. Panel B is a slice of the testing image, a real image of an X-ray tomography reconstruction. Panel D shows approximate ground truth of B created manually.

We tried to limit the size of the set of parameters controlling the simulated root and soil. First, the shape of the root systems is not chosen realistically (we found it was not mandatory), but the typical size of the object is chosen similar to the size of the real roots to be segmented. Also, like in the nominal conditions section, we tuned the first-order statistics (mean and standard deviation) of roots and soil given in Table 3 which corresponded to the contrast found in real acquisition conditions (Fig. 7, panel B). In addition, we tuned the second-order statistics of the simulated soil and roots on real data sets. These second-order statistics were controlled by mean of the algorithm described in Fig. 8. By operating this way, we obtained the very promising segmented images of Fig. 9 with a confusion matrix given in Table 4.

**Table 3:** Mean (*µ*) and standard deviation (*σ*) values for the roots and soil of the experiment with real root images. Images are coded on 8-bits. Training and testing images have the same statistics: simulated training images were created to resemble the testing ones.

| Image | Root $\mu$ | Root $\sigma$ | Soil $\mu$ | Soil $\sigma$ |
| --- | --- | --- | --- | --- |
| Statistics | 110 | 15 | 180 | 25 |

**Figure 8:**
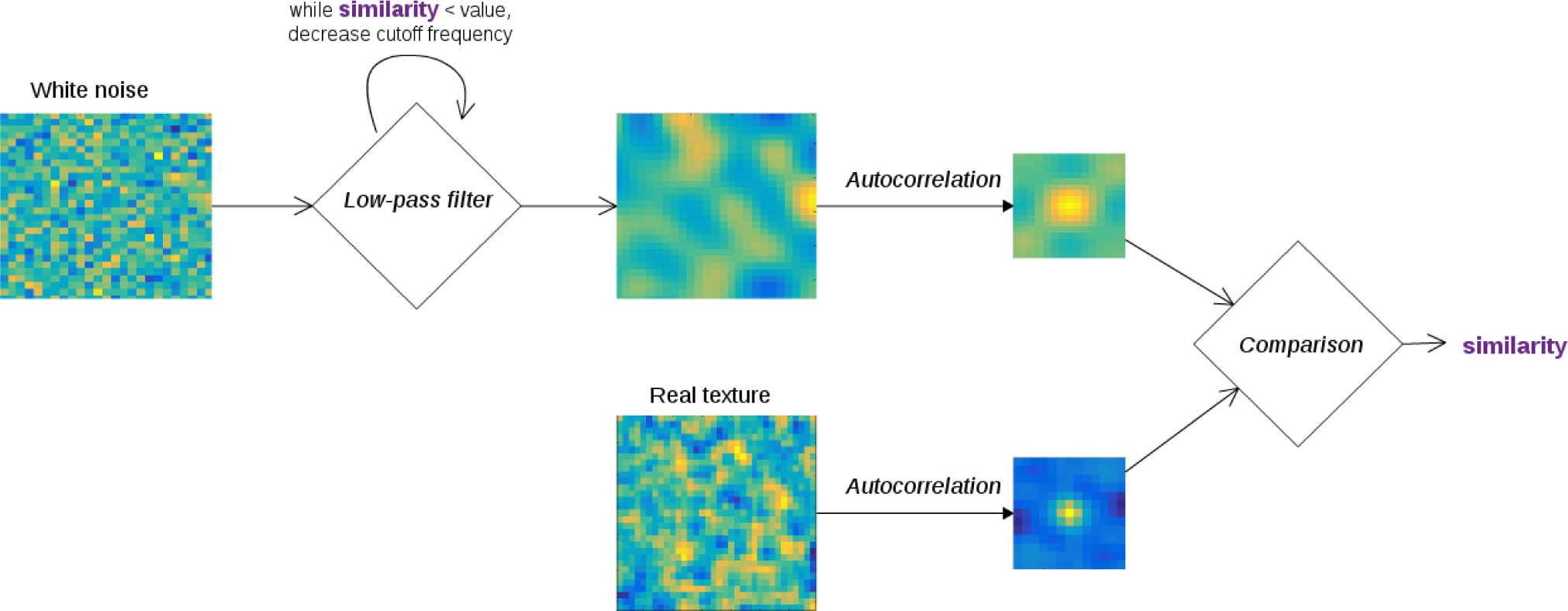
Pipeline for creating simulated texture with the same autocorrelation as real images. White noise is generated, then a low-pass filter is applied. This filter is realized by a multiplicative binary mask in the discrete cosine transform domain where low spatial frequencies are multiplied by 1 and the high spatial frequencies are multiplied by 0. The size of the mask acts as the cutoff frequency of this filter. We start with a very high cutoff frequency (*i.e.* we don’t change much the white noise). The autocorrelation matrix of the new image is then compared to the one of the real image, and the cutoff frequency is decreased if they are not similar enough, making the image “blurrier” and the autocorrelation spike wider.

**Figure 9:**
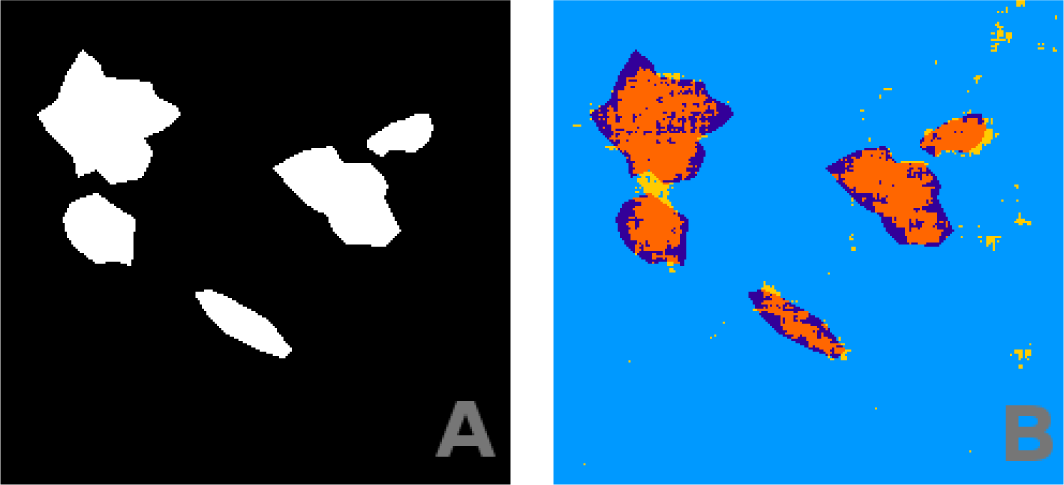
Panel A shows ground truth (manually estimated). Panel B shows the segmentation obtained from our ML algorithm, with parameters training set size = 1000, patch = 12. Color code is the same as in Fig. 3

**Table 4:** Confusion matrix of the results of Fig. 9. Total number of pixels was 49.248. QM=0.57

|  | Root pixel | Soil pixel |
| --- | --- | --- |
| Predicted root | 6.9% | 1.0% |
| Predicted soil | 3.7% | 88.4% |

### 4.1 Robustness

We have engineered the training data, generated from simulation, to enable our ML algorithm to yield the good segmentation result of Fig. 9. We found that the simulated training data must share some similar statistics with the real image for the segmentation to work. Specifically, it is sufficient to ensure the match of the first-order statistics (mean, standard deviation) and second-order statistics (autocorrelation) with the corresponding statistics of the image to be segmented. To test the robustness of this result, we have realized the same segmentation while changing one of these statistics in the training data, the other two staying similar to the testing data. Evolution of the performances can be found in Fig. 10.

**Figure 10:**
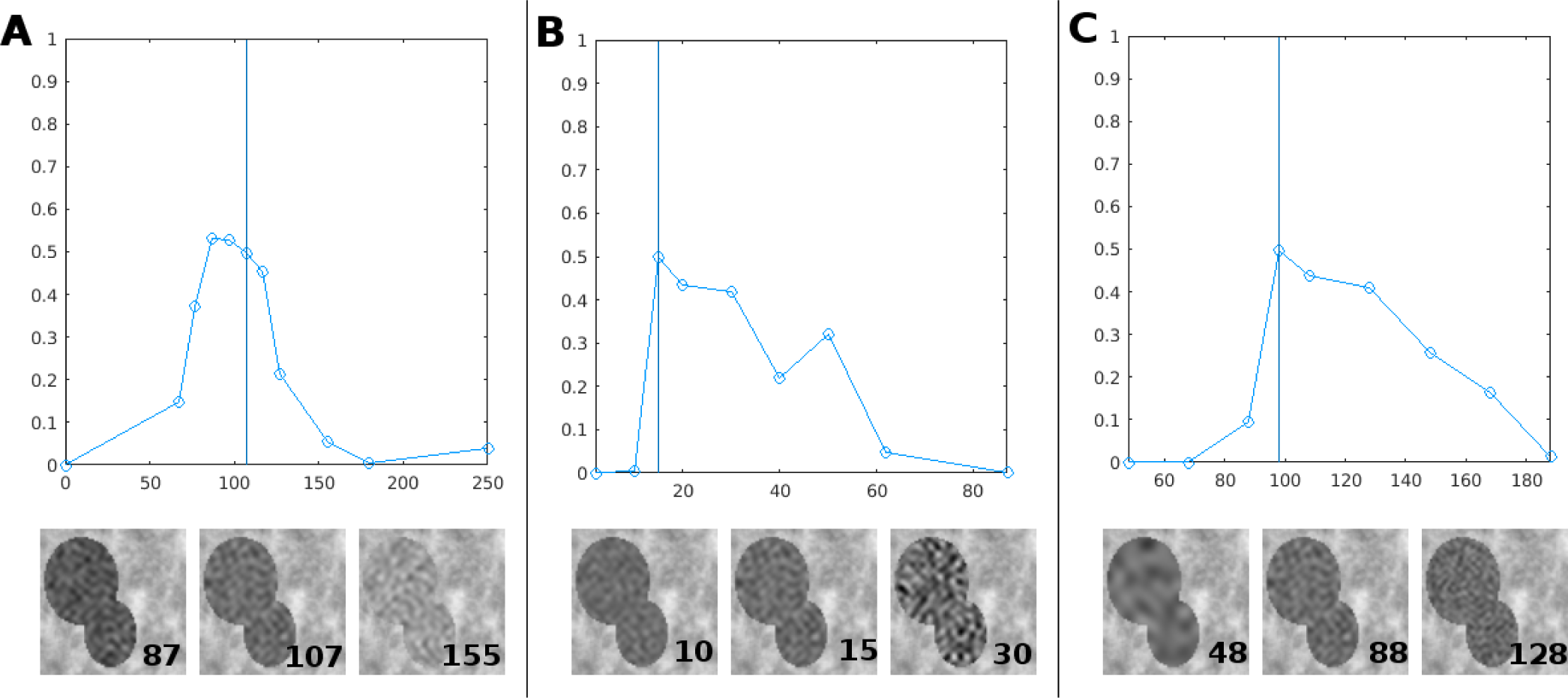
Panel A, top, shows evolution of QM when the root intensity mean varies from 0 to 255 in the training data (s.d. and autocorrelation staying the same as testing data). The vertical blue line is the value of root mean in the testing data (the one we used for our results of Fig. 9). Bottom part shows part of training images with different means as an illustration. Panels B and C show same results for standard deviation and autocorrelation. Autocorrelation was quantified as the value of the cutoff frequency of the filter during the simulated image’s creation.

As expected, segmentation is best when training data resembles testing data (vertical line in Fig. 10). However, segmentation remains good when statistics depart in a reasonable range around the optimal value. These ranges are visible in Fig. 10. For example, panel A shows that the mean of the root in the 80-120 range provides reasonably good results compared to the optimum case of 97 which corresponds exactly to the mean of the image to be segmented. This establishes conditions where it is possible to automatically produce efficient segmentation of soil and root in X-ray tomography. Such regimes of stability are found also in panels B and C of Fig. 10 for the standard deviation and the autocorrelation.

## 5. Conclusion

In this article, we have demonstrated the value of deep learning to address the difficult problem of soil-root segmentation from X-ray tomography images. This was obtained from the so-called transfer learning approach where a convolutional neural network is trained on a huge image data set, distinct from soil-root, for classification purposes to select a feature space which is then used to train an SVM on the soil-root segmentation problem. We demonstrated that such an approach gives very good results on simulated roots and on real roots even when the soil/root contrast is very low. We have discussed the robustness of the obtained results with respect to the size of the training data sets and the size of the patch used to classify each pixel. We illustrated the possibility to perform segmentation of real roots from training on purely synthetic soil and root. This was obtained in stationary conditions where both soil and root could be approximated by their first order and second order statistics.

These first uses of deep learning for the soil-root segmentation problem can serve as a reference to investigate more complex situations which are found in practice. The water density of root may not be constant along the root systems. The soil because of gravity, is often not found to present the same compactness along the vertical axis. It would therefore be important to push forward the investigation initiated in this article in the direction of non-stationarity of the gray levels in the root and in the soil. Also as discussed in section 3 of this article, this work opens many perspectives of optimization in the selection of the convolutional neural network, the extension of the patches in 3D and the post processing of the classified images.

It is important to underline again that the use of simulated data, which offers the possibility to generate unlimited data sets and which enables the control of all parameters of the data set, was specially useful to establish conditions on which transfer learning can be expected to give good results with real soil/root segmentation. Also, thanks again to the use of simulated data which creates annotated ground truth, our approach could serve to present a comparison with classic image analysis methods for soil/root segmentation (for instance Mairhofer et al. (2012)) or other deep learning based algorithms. However, transfer learning is a generic approach which can be applied whenever a classification task is targeted. It could thus also be applied to other computer vision problem for the plant sciences which can be expressed in classification terms such as the detection of pathogens and healthy tissue Neethirajan et al. (2006); Sankaran et al. (2010), the segmentation of shoot from background Chéné et al. (2012) or the classification of species Gwo et al. (2013) to cite only a few recent issues in plant phenotyping. This calls for the generation of annotated data sets or the production of simulators so as to establish in controlled conditions the typical performances to be expected with deep learning applied in plant sciences in the same way as initiated in this article.

## Acknowledgments

This work received supports from the French Government supervised by the Agence Nationale de la Recherche in the framework of the program Investissements d’Avenir under reference ANR-11-BTBR-0007 (AKER program). Richard SCHIELEIN acknowledges support from the EU COST action ‘The quest for tolerant varieties: phenotyping at plant and cellular level (FA1306)’.

